# Viral persistence and antiretroviral therapy shape systemic immune aging in treated HIV infection

**DOI:** 10.64898/2026.04.01.715809

**Authors:** Yubo Zhang, Vasiliki Matzaraki, Adriana Navas, Nadira Vadaq, Marc Blaauw, Willem Vos, Albert Groenendijk, Louise van Eekeren, Janneke Stalenhoef, Anca-Lelia Riza, Ioana Streata, Vinod Kumar, Collins K. Boahen, Marvin Berrevoets, Casper Rokx, Mareva Delporte, Leo A.B. Joosten, Cheng-Jian Xu, Yang Li, Linos Vandekerckhove, Andre van der Ven, Mihai G. Netea

## Abstract

Chronic infections can reshape immune system homeostasis, yet how persistent viral infections influence immune aging remains poorly understood. People living with HIV provide a unique model to investigate how long-term viral persistence affects immune aging despite effective antiretroviral therapy. Here, we characterize immune aging by integrating plasma proteomics with epigenetic and transcriptional profiles of circulating immune cells across large cohorts of treated individuals with HIV. We find that immune aging is markedly accelerated compared with healthy individuals and parallels established DNA methylation–based aging clocks. Accelerated immune aging is strongly associated with signatures of immunosenescence and correlates with the size of the latent HIV reservoir, suggesting a persistent imprint of viral persistence on immune aging trajectories. Notably, exposure to specific antiretroviral agents, particularly nucleoside reverse transcriptase inhibitors, is associated with reduced immune aging and suppression of age-associated immune gene programs. Together, these findings identify chronic viral persistence as a driver of systemic immune aging and indicate that antiretroviral therapy can partially modulate immune aging programs.

## Introduction

With early administration of combination antiretroviral therapy (cART), people with HIV (PWH) can now achieve life expectancies approaching those of uninfected individuals. Nevertheless, accumulating evidence indicates that PWH experience pronounced premature aging and an increased burden of age-associated comorbidities, including cardiovascular disease, metabolic disorders, and neurocognitive decline, even under effective viral suppression and especially when ART administration has been delayed. Chronic immune activation and inflammation have been proposed as central drivers of these processes, positioning immune dysfunction as a key mediator linking HIV infection to accelerated aging (*1–4*). Indeed, HIV-associated inflammation persists despite cART and has been strongly implicated in the development of non-AIDS–related comorbidities (*5*, *6*).

Advances in aging research have led to the development of biological clocks that quantify aging beyond chronological time. Epigenetic clocks based on DNA methylation patterns are among the most well-validated biomarkers of biological aging and robustly predict morbidity, mortality, and health span (*7–9*). In parallel, plasma proteomic signatures have emerged as powerful indicators of biological age, capturing systemic changes across the lifespan and providing insight into aging-related physiological decline (*10*). More recently, immune composition–based approaches, such as the IMM-AGE framework, have leveraged immune cell distributions to estimate immune aging and link immune system remodeling to clinical outcomes (*11*).

Immune aging is characterized by hallmark features including loss of naïve lymphocyte populations, expansion of differentiated effector and exhausted cells, and heightened basal inflammation, collectively contributing to impaired immune responsiveness (*12*, *13*). In the context of a treated HIV infection, these processes are further shaped by persistent viral reservoirs and ongoing immune activation. HIV reservoirs have been shown to induce sustained immune dysregulation even during suppressive therapy, thereby potentially accelerating immune aging trajectories (*14*, *15*). Notably, elite controllers—individuals who maintain viral suppression in the absence of cART—offer a unique model for dissecting the relationship between viral persistence, immune activation, and aging (*16*).

In our previous work, we integrated plasma proteomics and DNA methylation data to quantify systemic and organ-specific biological aging in PWH, demonstrating that chronic HIV infection and antiretroviral therapy exert opposing effects on aging trajectories across multiple organ systems (*17*). These findings raised the possibility that the immune system—both as a primary target of HIV infection and as a central regulator of systemic inflammation—may represent a critical axis through which HIV influences biological aging.

Building on this foundation, the present study focuses specifically on the biological aging of the immune system in PWH. We hypothesized that immune aging is accelerated in PWH that are virally suppressed relative to healthy individuals and that this process may be modulated by specific antiretroviral regimens. To test this hypothesis, we identified 139 immune-related plasma proteins using the Genotype-Tissue Expression (GTEx) resource and trained a proteomic immune-aging model in healthy individuals from the 200FG cohort of the Human Functional Genomics Project. We then applied this model to two cohorts of PWH (200HIV and 2000HIV) to derive immune aging scores. These scores were systematically compared with DNA methylation–based aging measures and immune composition–based aging estimates, and their associations with clinical characteristics, HIV disease parameters, comorbidities, HIV reservoir size, and both current and cumulative antiretroviral exposure were evaluated.

By integrating transcriptomic data from an independent healthy cohort with those from PWH, we identified age-associated gene programs modulated by antiretroviral therapy, providing mechanistic insight into immune aging in chronic HIV infection (Figure 1).

**Figure 1.**
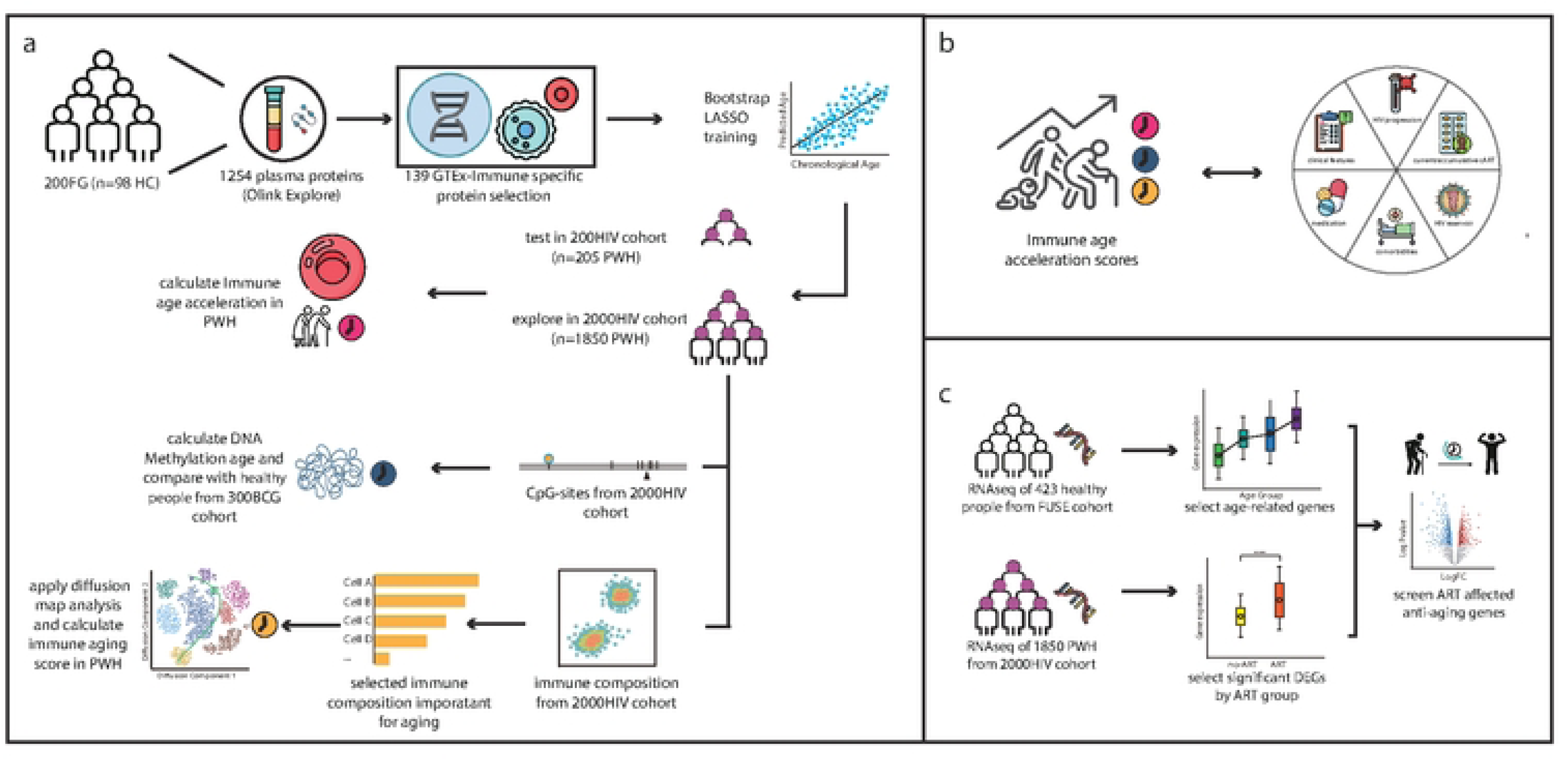
Overview of the study design and analytical framework for immune aging in people with HIV. (a) Proteomic immune aging clock construction and validation. Plasma proteomic profiles measured by Olink Explore were obtained from healthy individuals in the 200FG cohort (n = 98). Immune-specific proteins (n = 139) were selected based on GTEx expression data and used to train a proteomic immune aging model using bootstrap LASSO regression. The trained model was subsequently applied to two independent cohorts of people with HIV (PWH), the 200HIV cohort (n = 205) for validation and the 2000HIV cohort (n = 1,850) for large-scale exploratory analyses, to estimate immune age and immune age acceleration. DNA methylation age and immune composition–based aging scores were additionally calculated for comparative analyses. (b) Multi-layer characterization of immune aging in PWH. Immune age acceleration scores were systematically associated with clinical characteristics, HIV disease progression parameters, antiretroviral therapy (ART) exposure (current and cumulative), comorbidities, medication use, and HIV reservoir size (c) Identification of ART-modulated age-associated immune transcripts. Bulk RNA-seq data from healthy individuals in the FUSE cohort (n = 423) were used to define age-associated genes. These signatures were integrated with bulk transcriptomic profiles from PWH in the 2000HIV cohort (n = 1,850) to identify genes whose age-associated expression patterns were modulated by ART exposure, highlighting molecular pathways linking antiretroviral treatment to immune aging.

## Results

### Plasma proteomics–based immune age and DNA methylation age reveal immune age acceleration in PWH

To construct a proteomics-based immune aging model, we first identified 139 immune-specific proteins from 1,254 plasma proteins quantified by the Olink platform. These proteins exhibited at least a two-fold higher expression in immune-related tissues compared with other major organs profiled in the GTEx dataset (Fig. 2a). Principal component analysis based on normalized expression of these immune-specific proteins revealed no evident outlier separation among healthy participants (200FG) and PWH from the 200HIV and 2000HIV cohorts (Fig. 2b), indicating comparability across cohorts.

**Figure 2.**
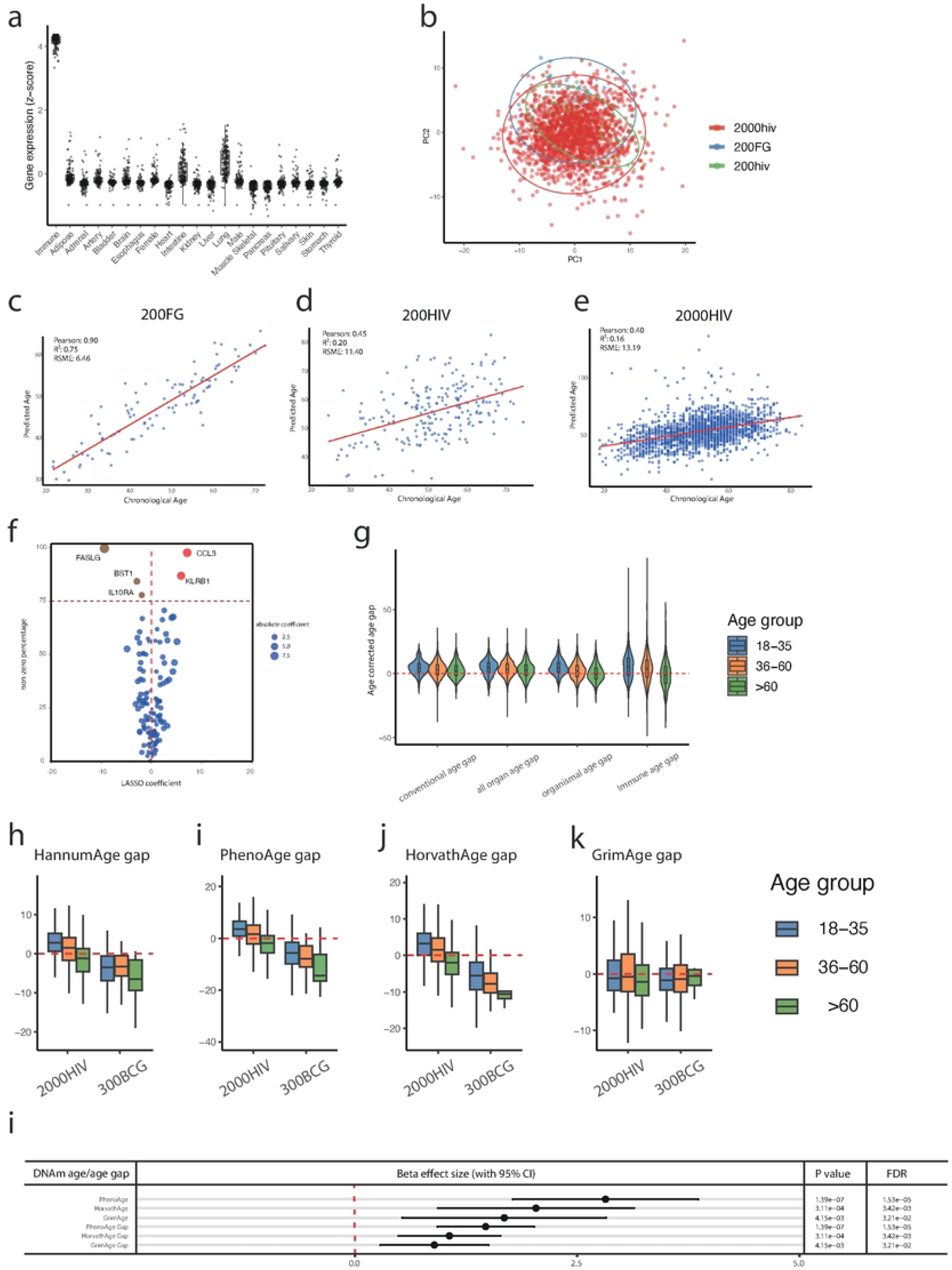
Proteomic immune aging and DNA methylation aging reveal immune age acceleration in PWH. (a) Identification of immune-specific proteins based on GTEx transcriptomic data. Proteins exhibiting at least a two-fold higher expression in immune-related tissues relative to other major organs were classified as immune-specific. (b) Principal component analysis of normalized expression of immune-specific proteins across healthy individuals (200FG) and PWH cohorts (200HIV and 2000HIV). (c) Performance of the proteomic immune aging clock in the 200FG training cohort, showing the relationship between predicted immune age and chronological age. (d,e) Application of the immune aging clock to PWH in the 200HIV (d) and 2000HIV (e) cohorts. (f) LASSO regression coefficients and selection stability of immune-specific proteins contributing to immune age prediction. (g) Age-corrected immune age gaps across age strata in PWH, compared with previously reported systemic aging measures. (h–k) Residual DNA methylation age gaps (HannumAge, PhenoAge, HorvathAge, and GrimAge) in PWH from the 2000HIV cohort and healthy individuals from the 300BCG cohort, stratified by age group. (l) Associations between proteomic immune age acceleration and DNA methylation age measures and age gaps in PWH. Effect sizes (β), 95% confidence intervals, P values, and false discovery rates are shown.

Using these immune-specific proteins, we trained an immune aging clock in healthy individuals from the 200FG cohort (n = 98) using bootstrap-aggregated LASSO regression (Fig. 1a). Across 500 bootstrap iterations, predicted immune age showed a strong correlation with chronological age (Pearson’s r = 0.90), with a coefficient of determination (R²) of 0.75 and a root mean squared error (RMSE) of 6.46 years (Fig. 2c). Out-of-bag (OOB) validation yielded an R² of 0.33 and an RMSE of 10.56 years (Table S2), supporting the robustness and partial generalizability of the model. Within the trained model, CCL3 and KLRB1 showed the strongest positive contributions to immune age prediction, whereas FASLG, BST1, and IL10RA showed the strongest negative contributions (Fig. 2f).

We next applied this immune aging model to PWH in the 200HIV and 2000HIV cohorts. In both cohorts, predicted immune age remained moderately correlated with chronological age (r = 0.45 and r = 0.40, respectively; Fig. 2d,e). Notably, consistent with previously reported systemic aging clocks, including conventional age, all-organ age, and organismal age, the immune age scores exhibited significant acceleration in PWH aged 18–35 years and 36–60 years. In contrast, no immune age acceleration was observed among individuals older than 60 years, consistent with patterns reported for systemic proteomic aging. (Fig. 2g).

To contextualize proteomic immune aging within established biological aging frameworks, we estimated four DNA methylation–based age metrics (HannumAge, PhenoAge, HorvathAge, and GrimAge) using methylation data from the 2000HIV cohort (n = 1,850) and healthy individuals from the 300BCG cohort (n = 286). Residual age gap analyses revealed significant age acceleration in PWH for HannumAge, PhenoAge, and HorvathAge, particularly in younger age groups, mirroring patterns observed for immune and systemic proteomic aging (Fig. 2h–j). In contrast, GrimAge did not differ significantly between PWH and healthy individuals (Fig. 2k).

Across PWH, proteomic immune age acceleration was significantly associated with multiple DNAm age measures, including PhenoAge, HorvathAge, and GrimAge, as well as their corresponding age gaps, with PhenoAge showing the largest effect size (β = 2.82; Fig. 2l). These findings are consistent with our previous observations of DNA methylation age acceleration in PWH and support the presence of coordinated immune and epigenetic aging processes in people with HIV with long-term cART exposure.

### Clinical and immunological correlates of immune aging in PWH

To characterize the clinical relevance of immune aging in PWH, we systematically assessed associations between the corrected proteomic immune age gap and a broad range of clinical parameters in the 2000HIV cohort using multivariable linear models (Fig. 3a). Proteome immune age scores showed significant positive associations with markers of renal and hepatic function, including serum creatinine (CREAT), alanine aminotransferase (ALAT), body mass index (BMI), body weight, and liver stiffness measurement (LSM), while exhibiting a negative association with estimated glomerular filtration rate (eGFR). Because dolutegravir use is known to influence circulating creatinine levels, we performed additional analyses adjusting for current dolutegravir exposure. The association between immune age acceleration and serum creatinine levels remained robust after adjustment (β = 1.35, P = 5.56 × 10⁻⁷; Fig. S1), indicating that this relationship is not solely attributable to dolutegravir-related creatinine changes. Together, these findings link accelerated immune aging to impaired liver and kidney function in PWH.

**Figure 3.**
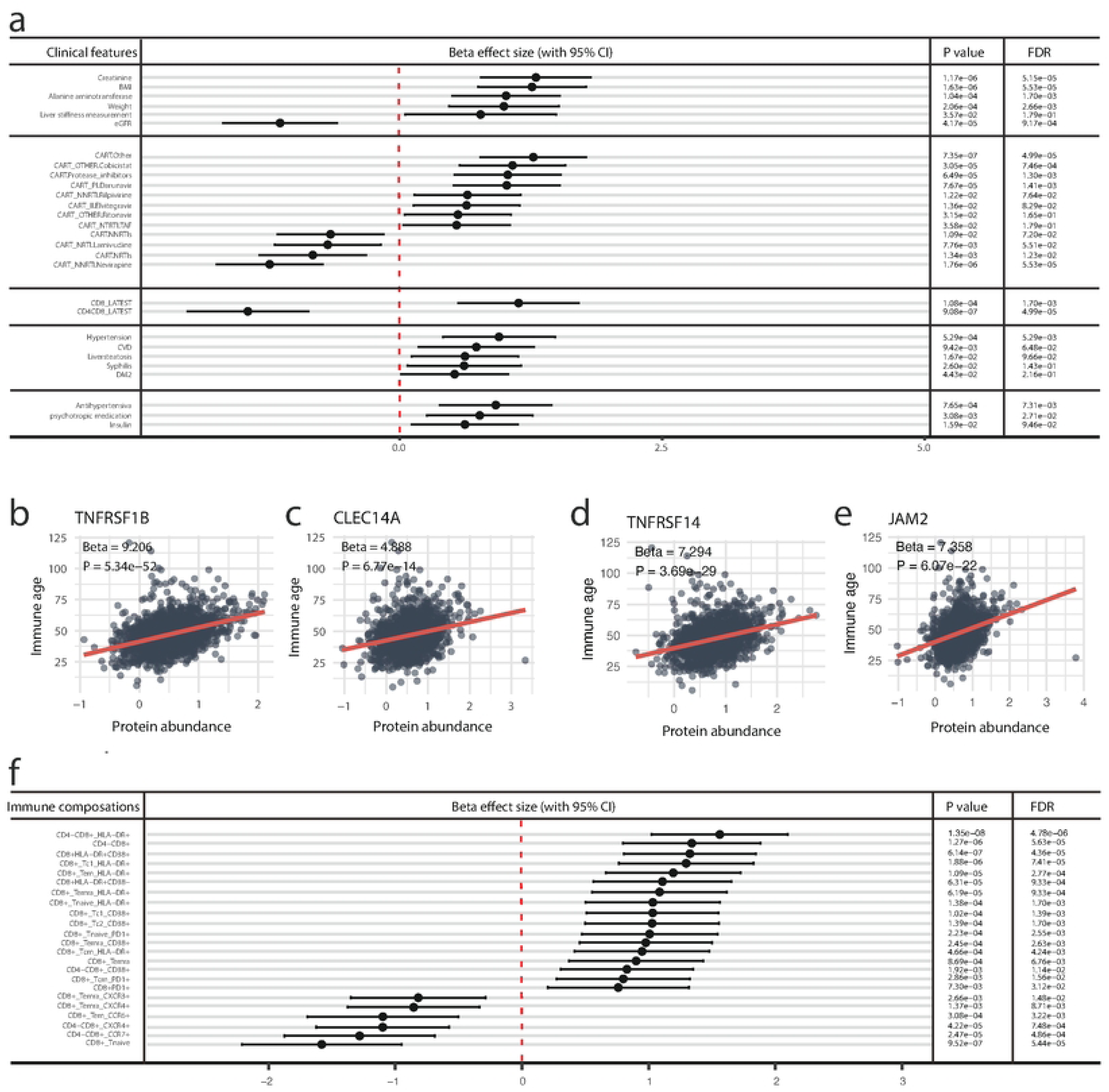
Clinical and immunological correlates of proteomic immune aging in PWH. (a) Associations between corrected proteomic immune age gap and clinical features in the 2000HIV cohort, assessed using multivariable linear models. Effect sizes (β) with 95% confidence intervals, P values, and false discovery rates (FDR) are shown. (b–e) Representative associations between immune age and HIV-associated heart failure biomarkers with high reported hazard ratios, including TNFRSF1B, CLEC14A, TNFRSF14, and JAM2. (f) Meta-analysis of associations between circulating immune cell subsets and immune age acceleration, highlighting negative associations with naïve T-cell populations and positive associations with differentiated, activated, or exhausted T-cell subsets.

Distinct associations were observed between immune aging and different antiretroviral therapy (ART) drug classes. Current exposure to reverse transcriptase inhibitors—particularly the nucleoside reverse transcriptase inhibitor lamivudine (3TC) and the non-nucleoside reverse transcriptase inhibitor nevirapine (NVP)—was negatively associated with immune age acceleration, whereas protease inhibitors and other ART classes showed positive associations. No significant association was detected for integrase strand transfer inhibitors (INSTIs). These patterns parallel previously reported effects of ART on systemic aging markers (*18*), suggesting that specific ART agents may exert a decelerating effect on immune aging, whereas others may contribute to immune age acceleration.

Consistent with hallmarks of immune senescence, immune age acceleration was positively associated with CD8⁺ T-cell counts and negatively associated with the CD4⁺/CD8⁺ T-cell ratio, while no significant association was observed with absolute CD4⁺ T-cell counts alone (Fig. S2). In addition, immune aging was significantly associated with multiple HIV-related comorbidities, including hypertension, cardiovascular disease, liver steatosis, a history of syphilis infection, and type 2 diabetes mellitus, as well as with the use of antihypertensive medications, insulin, and psychiatric drugs. In contrast, no significant associations were observed between proteome -based immune age scores and malignancies, although the prevalence of malignancy was lower than other diseases and one has to be cautious regarding the power to be able to identify such associations.

Given the elevated burden of cardiovascular disease among PWH, we next examined the relationship between immune aging and 26 previously reported proteins associated with left atrial remodeling and incident clinical heart failure in people with HIV (*19*). Strikingly, immune age was significantly associated with 25 of the 26 markers, with the exception of NT-proBNP (Table S3). Notably, biomarkers with the highest reported hazard ratios for incident heart failure—including TNFRSF1B, CLEC14A, TNFRSF14, and JAM2—showed particularly strong associations with immune age acceleration (Fig. 3b–e). These results underscore a close link between immune aging and cardiovascular risk in PWH.

To further dissect the immunological basis of immune aging, we analyzed 356 circulating immune cell subsets quantified by high-dimensional flow cytometry in 1,296 individuals from the 2000HIV cohort. Meta-analyses revealed that naïve CD4⁺ and CD8⁺ T cells numbers— canonical indicators of immunological youth—were negatively associated with immune age acceleration. Similarly, TCRγδ2 (TCRVδ2) cell abundance was inversely associated with immune aging (Fig. 3f; Figs. S3 and S4). In contrast, effector memory (T_EM) and effector memory RA (T_EMRA) CD8⁺ T cells exhibited strong positive associations with immune age acceleration. Moreover, CD4⁺ T cells, CD8⁺ T cells, and TCRγδ cells expressing activation or exhaustion markers, including HLA-DR, CD38, and PD-1, showed consistent positive associations with immune aging. Notably, even subsets of phenotypically naïve CD8⁺ T cells expressing HLA-DR or PD-1 were positively associated with immune aging, highlighting widespread activation and differentiation shifts across the T-cell compartment. Collectively, these findings demonstrate that proteomic immune aging in PWH captures clinically meaningful, ART-modifiable immunosenescence processes that are reflected in both immune cell composition and cardiovascular risk–associated molecular signatures.

### Immune composition–based aging and the imprint of viral persistence

We next applied bootstrap-based LASSO feature selection to identify immune cell subsets that most strongly contributed to chronological age in PWH (Fig. 4a; Table S4). Thirteen immune cell populations consistently retained non-zero coefficients across 500 bootstrap iterations and were subsequently used to derive an immune composition–based aging metric. Diffusion pseudotime analysis of these cell populations revealed a dominant and continuous immunological trajectory across the cohort (Fig. 4b), and scaled pseudotime values were interpreted as relative immune composition age. Immune composition age acceleration was associated with CD8⁺ T-cell counts and selected comorbidities, including central nervous system disorders, thyroid dysfunction, and psychiatric conditions (Fig.4c). However, compared with the immune composition–based metric, the proteomic immune aging score demonstrated a broader spectrum of significant associations and stronger statistical signals across clinical and immunological features.

**Figure 4.**
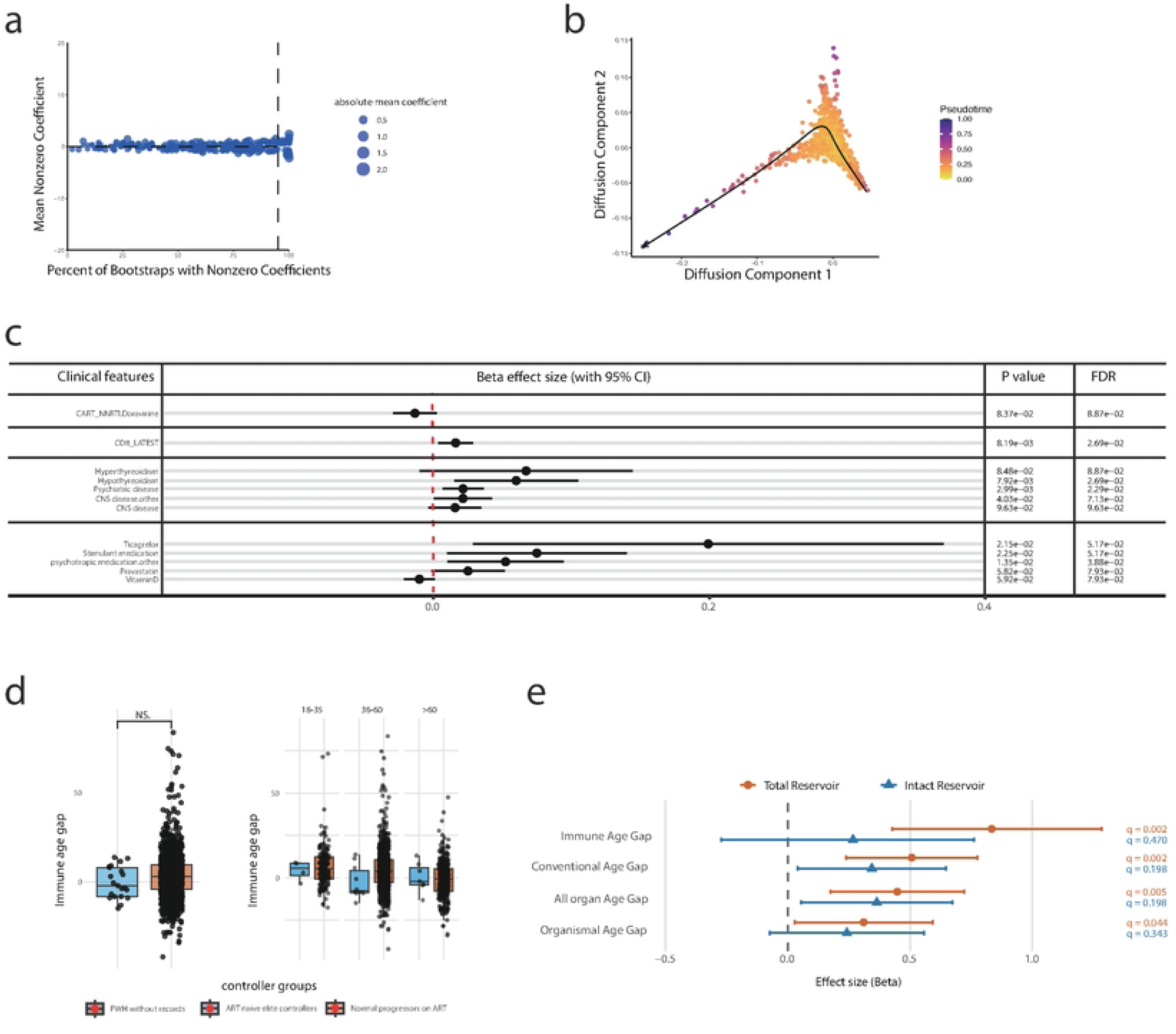
Immune composition–based aging trajectories and the relationship to HIV reservoir size. (a) Bootstrap LASSO feature selection identifying immune cell subsets contributing to chronological age, showing the proportion of bootstrap iterations with non-zero coefficients and mean effect sizes. (b) Diffusion pseudotime analysis based on selected immune cell populations, illustrating a continuous immune composition trajectory across PWH. (c) Meta-analysis of associations between immune composition age acceleration and clinical features, demonstrating that the proteomic immune aging metric yields a greater number of significant associations and stronger statistical signals. (d) Immune age residuals in elite controllers and normal progressors stratified by age group (18–35, 36–60, >60 years). (e) Comparison of associations between HIV reservoir size and different biological aging metrics, including conventional age gap, all-organ age gap, organismal age gap, and proteomic immune age gap. Effect sizes and false discovery rates are shown.

We further quantified immune age across the full dataset, including all 21 elite controllers, and examined differences between elite controllers and normal progressors both overall and across age strata (18–35, 36–60, and >60 years; Fig. 4d). Notably, elite controllers— particularly in the middle-aged group—displayed a trend toward immune age deceleration compared with normal progressors, consistent with partial preservation of immune youth in the absence of sustained viral replication. To define the contribution of viral persistence to immune aging, we next examined associations between HIV reservoir size, quantified by analyzing total and intact HIV DNA in circulating CD4 T cells, and immune aging across molecular layers. Proteomic immune age acceleration showed a significant association with total HIV DNA levels, exceeding the strength of association observed for conventional age gap, all-organ age gap, or organismal age gap derived from previous systemic aging analyses (Fig. 4e).

Collectively, these results demonstrate that proteomic immune aging captures systemic and clinically relevant features of immune senescence in people with HIV that are not fully reflected by circulating immune cell composition alone. In line with prior systemic aging analyses, viral reservoir burden preferentially imprints on proteome-based immune aging signatures, highlighting the added value of plasma proteomics for quantifying HIV-associated immune aging.

### Transcriptional signatures and antiretroviral modulation of immune aging

Given the observed association between HIV reservoir size and proteomic immune aging, we next sought to determine whether antiretroviral drugs modulate immune aging at the transcriptional level. We first characterized age-associated transcriptional programs in healthy individuals using bulk PBMC RNA-sequencing data from 423 participants in the FUSE cohort. Spearman correlation analysis between gene expression levels and chronological age identified 1,299 age-associated genes showing moderate-to-strong correlations with chronological age (correlation coefficient > 0.3 or < −0.3; Fig. 5a, Table S5). Among these, 33 genes exhibited particularly strong age associations (|ρ| > 0.5; Fig. S5), indicating robust transcriptional responses to aging. Functional enrichment analyses revealed that genes negatively correlated with age were predominantly enriched in pathways related to lymphocyte differentiation and T cell development (Fig. 5b), whereas genes positively correlated with age were mainly associated with myeloid cell functions, immune activation, and inflammatory responses (Fig. 5c).

**Figure 5.**
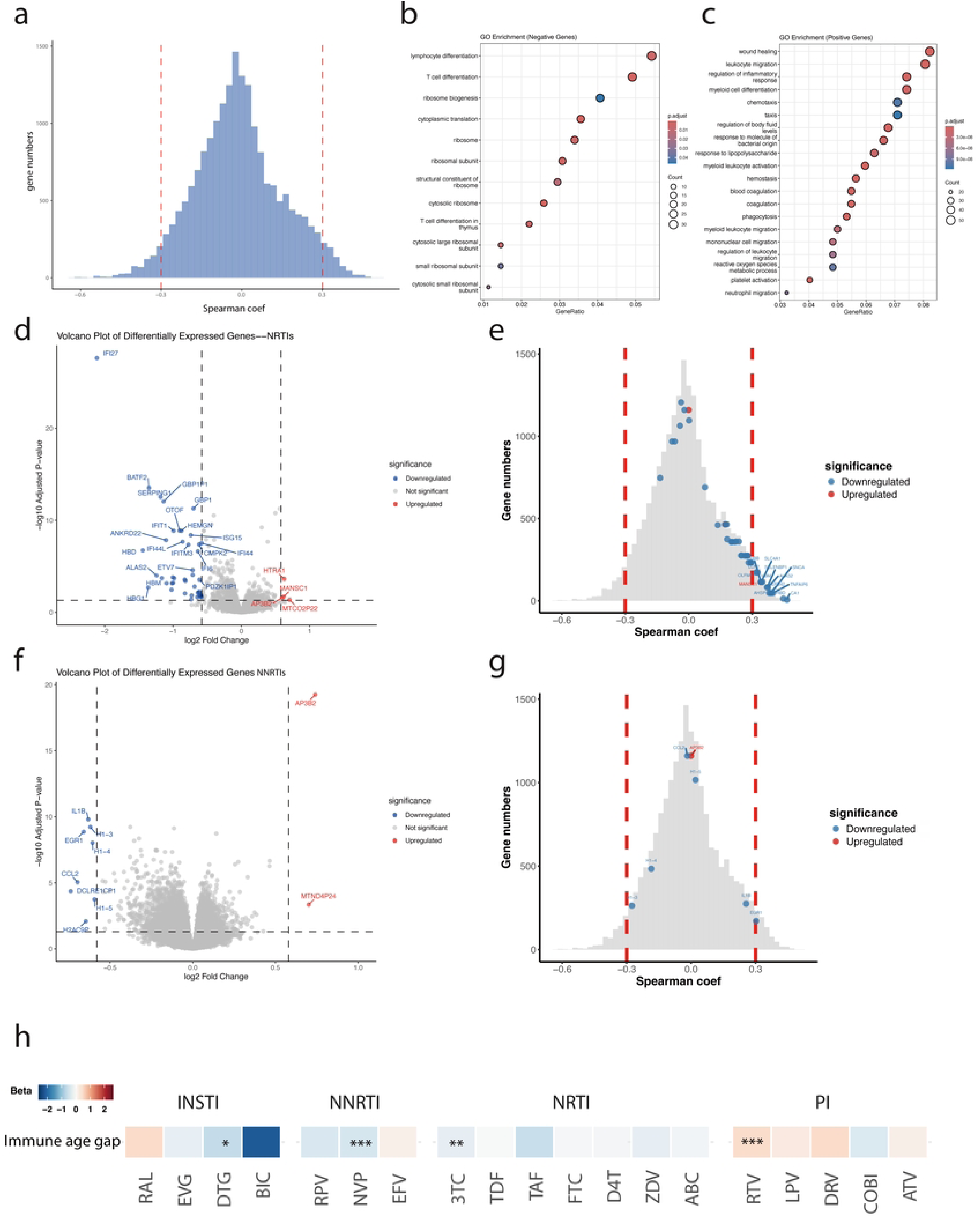
Antiretroviral therapy modulates age-associated immune transcriptional programs. (a) Distribution of Spearman correlation coefficients between gene expression levels and chronological age in healthy individuals from the FUSE cohort, identifying age-associated genes. (b) Gene Ontology enrichment analysis of genes positively associated with age, highlighting pathways related to lymphocyte differentiation and T-cell development. (c) Gene Ontology enrichment analysis of genes negatively associated with age, primarily enriched in myeloid cell functions and inflammatory processes. (d) Volcano plot showing differential gene expression associated with current NRTI use in PWH. (e) Overlay of differentially expressed genes in NRTI users on the age-correlation distribution, demonstrating preferential downregulation of positively age-associated genes. (f) Volcano plot showing differential gene expression associated with current NNRTI use in PWH. (g) Distribution of age-associated genes modulated by NNRTI use, highlighting overlaps with positively age-associated transcripts. (h) Associations between immune age gap and cumulative exposure to major antiretroviral drug classes and individual agents, adjusted for co-administration effects.

We next investigated whether exposure to classes of ART agents modulate these aging-related transcriptional patterns in PWH. Using bulk PBMC RNA-seq data from the 2000HIV cohort, we performed differential expression analyses comparing individuals currently receiving nucleoside reverse transcriptase inhibitors (NRTIs; abacavir, lamivudine, emtricitabine; n = 1,758) or non-nucleoside reverse transcriptase inhibitors (NNRTIs; efavirenz, etravirine, nevirapine, rilpivirine, doravirine; n = 727) with individuals not receiving these drug classes (n = 92 and n = 1,123, respectively). Current NRTI use was associated 47 differentially expressed genes, with widespread downregulation of transcripts, including IFI27, BATF2, SERPING1, and GBP1P1, whereas only a limited number of genes—such as HTRA1, MANSC1, AP3B2, and MTCO2P22—were upregulated (Fig. 5d). In contrast, NNRTI use was associated with fewer transcriptional changes, with significant downregulation of IL1B and EGR1 and upregulation of AP3B2 (Fig. 5f).

Strikingly, although current NRTI use was associated with a selective set of differentially expressed genes (n = 47), the majority of downregulated genes overlapped with genes positively associated with chronological aging (Table S6). Specifically, multiple aging-associated genes with Spearman correlation coefficients > 0.3—including CA1, TNFAIP6, HBD, SNCA, EPB42, ALAS2, HBM, SELENBP1, SLC4A1, AHSP, LCN2, HBB, and OLFM4— were significantly downregulated in NRTI-treated individuals (Fig. 5e). These findings suggest that NRTI use selectively suppresses age-associated immune transcriptional programs.

Although NNRTI-associated transcriptional changes did not reveal a consistent overall directionality, several positively age-associated genes—most notably IL1B and EGR1—were also significantly downregulated in individuals receiving NNRTIs (Fig. 5g), partially overlapped with aging-associated genes.

Beyond current drug exposure, we examined the relationship between cumulative exposure to ART agents and immune aging. Prolonged cumulative use of dolutegravir (DTG), nevirapine (NVP), and lamivudine (3TC) was significantly associated with reduced proteomic immune aging, whereas cumulative exposure to ritonavir (RTV) was associated with increased immune age acceleration. Because antiretroviral drugs are administered in combination regimens, we systematically evaluated all commonly used treatment combinations in the 2000HIV cohort and applied regression models adjusting for co-administration effects. After accounting for combination therapy, independent associations remained, indicating that NVP, 3TC, and tenofovir alafenamide (TAF) were each independently associated with reduced immune aging, whereas elvitegravir (EVG) and darunavir (DRV) were associated with accelerated immune aging (Table S7). Collectively, while these associations are observational and cannot establish causality, they indicate that cART—particularly NRTIs and NNRTIs—can modulate immune aging at the molecular level and may partially counteract aging-associated transcriptional programs in PWH.

## Discussion

Growing interest in organ-specific aging has highlighted the heterogeneity and multidimensional nature of biological aging across human tissues and systems. Building on our previous work that quantified systemic and organ-specific aging using plasma proteomics data from the 2000HIV cohort, the present study extends this framework to the immune system—one of the primary targets of HIV infection. Here, we developed a proteomics-based immune aging score derived from immune-specific plasma biomarkers and systematically evaluated its biological and clinical relevance in people with HIV (PWH).

Using this approach, we demonstrate that PWH exhibit marked premature immune aging, consistent with acceleration observed using established DNA methylation–based epigenetic clocks. Beyond chronological aging, immune age acceleration was strongly associated with a broad range of clinical, virological, and immunological features, including total (but not the intact) HIV reservoir size, cardiovascular risk markers, liver and renal abnormalities, and canonical signatures of T-cell and γδ T-cell immunosenescence. Notably, the total HIV reservoir size strongly related to immune age acceleration, in contrast to the intact proviral DNA which presents only a minor fraction of the reservoir size. Furthermore, immune aging quantified at the proteomics level captured these associations more robustly than immune composition–based aging measures, indicating that circulating inflammatory and functional immune states contribute to immune aging scores beyond changes in cell subset abundance.

Finally, both current and cumulative exposure to specific antiretroviral agents—particularly reverse transcriptase inhibitors—were associated with reduced immune age acceleration, and current NRTI use selectively suppressed age-associated immune transcriptional programs. Together, these findings position immune aging as an integrative and therapeutically modifiable dimension of chronic HIV-associated immunopathology.

Immune aging is increasingly recognized as a critical determinant of health span, given the central role of the immune system in antimicrobial defense, cancer surveillance, tissue homeostasis, and tissue repair. As such, accelerated immune aging contributes broadly to age-related morbidity and functional decline. Importantly, immune aging often predicts adverse health outcomes more accurately than chronological age, reflecting the cumulative impact of lifelong immune stimulation driven by chronic infections, low-grade inflammation, and metabolic stress. In this context, the proteomic immune aging score developed in our study appears to capture an integrative dimension of immune aging, showing stronger and more consistent associations with clinical comorbidities and immune dysfunction than immune composition–based aging metrics. These findings support the notion that immune aging is not solely determined by shifts in cell subset abundance, but also by systemic inflammatory and functional immune states.

Two canonical hallmarks of immune aging—progressive loss of immune responsiveness and persistent low-grade inflammation—were evident in our analyses. Higher immune aging scores were associated with impaired T-cell functional states alongside elevated levels of inflammatory and cardiometabolic biomarkers, consistent with established features of immunosenescence and inflammaging. Notably, associations with antiretroviral therapy exposure further suggest that immune aging trajectories may be modifiable. Current and cumulative use of nucleoside reverse transcriptase inhibitors (including abacavir, lamivudine, and emtricitabine) were associated with reduced immune aging, accompanied by downregulation of age-associated immune transcriptional programs, indicating potential drug-class–specific effects on immune aging pathways. Apart from that, as the total HIV reservoir size is strongly associated with immune aging, the effect of reservoir reducing interventions on immune aging should be studied.

Several limitations should be considered when interpreting these findings. First, differences in proteomic coverage between cohorts limited the number of overlapping proteins available for immune aging model training, and the relatively modest size of the healthy training cohort may constrain detection of inter-individual heterogeneity. Second, the cross-sectional design precludes direct causal inference and limits assessment of temporal dynamics; longitudinal sampling will be essential to establish trajectories of immune aging over time. Third, the study population was predominantly male and of European ancestry, which may limit generalizability despite statistical adjustment for sex and ancestry. Future studies in more diverse and longitudinal cohorts will be important to validate and extend these observations.

In summary, by developing a proteomics-based immune aging score and integrating clinical, molecular data, we provide a comprehensive framework to quantify immune aging in people with HIV. Our findings reveal pronounced immune age acceleration with close concordance to DNA methylation–based aging measures, robust associations with immunosenescence signatures, comorbidities, and viral burden, and evidence that immune aging is partially modifiable by antiretroviral therapy. Collectively, this work establishes immune aging as an integrative and therapeutically tractable dimension of chronic HIV-associated immune dysregulation and provides broader insights into immune aging under conditions of sustained inflammatory stress.

## Materials and Methods

### Human cohorts

Two independent cohorts of people with HIV (PWH) that are virally suppressed—the 200HIV cohort (n = 205) and the 2000HIV cohort (n = 1,850, NTC03994835)—were included in this study. As a reference population without HIV, we analyzed the 200FG cohort (n = 98), enrolled as part of the Human Functional Genomics Project (HFGP). In addition, two healthy cohorts, the FUSE cohort (n = 423) and the 300BCG cohort (n = 286), were used for specific downstream analyses. Comprehensive descriptions of the healthy and PWH cohorts have been published previously (*20–22*).

For the present study, we integrated plasma proteomic profiles from 98 participants in the 200FG cohort, 205 participants in the 200HIV cohort, and 1,850 participants in the 2000HIV cohort. We further incorporated DNA methylation data, immune cell composition estimates, single-cell RNA sequencing (scRNA-seq) data, and detailed clinical parameters to enable multidimensional characterization across cohorts. cART is commonly prescribed as combination of three antiretroviral drugs: current and previous administration of all antiretroviral therapy (ART) agents were recorded, so cumulative exposure to specific antiretroviral agents and the associated drug class, could be calculated. Clinical summaries for the 200FG, 200HIV, and 2000HIV cohorts are provided in Table S8.

### Proteomic profiling of circulating plasma proteins

Circulating plasma proteins were quantified using the Olink proximity extension assay coupled to next-generation sequencing readout (Olink Proteomics AB, Uppsala, Sweden), as described previously (*23*). Protein abundances were reported as Normalized Protein eXpression (NPX) values on a log₂ scale.

Plasma samples from the 2000HIV and 200FG cohorts were profiled across three analytical batches using Olink’s Explore platforms. The first batch (n = 692) was assayed with the Olink® Explore 1536 panel (1,472 proteins spanning inflammation, oncology, cardiometabolic, and neurology pathways). The second batch (n = 692) used the Explore Expansion 1536 panel (1,472 proteins with extended content). The third batch was measured with the Olink® Explore 3072 platform (around 3,500 proteins across eight panels). For the present analyses, data from the first and third batches (1,500 proteins) were included.

Batch effects were corrected using bridging normalization as described previously (*23*). Briefly, median differences between bridging samples were computed and applied to harmonize NPX values across batches, with detection thresholds recalibrated using the same correction factors.

Following normalization, standard sample- and protein-level quality control was performed. Technical replicates (IL-6, TNF, CXCL8) included within the assay exhibited high reproducibility (Spearman r > 0.9). Proteins detected below the lower limit of detection (LOD) in ≥25% of samples were removed, yielding 1,306 proteins retained for the 2000HIV and 200FG datasets. Principal component analysis (PCA) was used to identify outlier samples (>4 SD from the mean on PC1 or PC2), resulting in 1,850 PWH samples and 98 healthy samples included in downstream analyses.

Proteomic profiling for the 200HIV cohort was performed using the Olink® Explore 1536 platform. The same normalization and QC procedures were applied, resulting in 1,254 high-quality proteins across 205 participants for final analyses.

### Immune cells count measurements

Peripheral blood samples from participants in the 2000HIV cohort were immunophenotyped using three dedicated flow cytometry panels on a 21-color, six-laser CytoFLEX LX instrument (Beckman Coulter), following previously established protocols (*24*). Instrument performance was monitored and standardized daily using CytoFLEX Daily QC and IR QC fluorospheres (Beckman Coulter, Cat. #B53230 and #C06147) and SPHERO™ Rainbow calibration particles (Spherotech Inc., Cat. #RCP-30-5A-6). Data acquisition was performed with CytExpert v2.3 (Beckman Coulter), and downstream analyses were conducted using conventional gating strategies implemented in Kaluza v2.1.2.

A total of 355 immune cell populations were defined, encompassing major innate, T-cell, and B-cell subsets. To assess functional perturbations, markers including HLA-DR, CD38, PD-1, PD-L1, CD40, CD307d, and CD81 were incorporated to evaluate activation, exhaustion, maturation, and intercellular communication states. The complete antibody panel has been described previously (*24*).

### Identification of immune-specific proteins

Normalized bulk RNA-seq expression data from the Genotype-Tissue Expression (GTEx) project (*25*) were used to identify immune-specific genes, as described previously (*23*).

Briefly, expression profiles from whole blood and spleen were combined, and genes exhibiting at least a two-fold higher expression in these immune-related tissues compared with any other organ were classified as immune-specific.

Immune-specific genes were subsequently mapped to proteins quantified in our Olink proteomic dataset. In total, 139 proteins corresponding to these genes were identified, representing ∼11% of all proteins measured. These immune-specific proteins were used to construct the proteomic immune aging model.

### Immune-specific aging clock training and immune age gap calculation

A bootstrap LASSO regression framework was used to train the immune-specific aging model, using expression levels of immune-specific proteins in the 200FG cohort as input features. For each bootstrap iteration (n = 500), a LASSO model was fit and used to generate age predictions for the training samples; the final predicted age for each individual was computed as the average across all bootstrap iterations. Model performance was evaluated by the mean prediction accuracy across the 500 bootstrap-derived models (Table S1) and by corresponding out-of-bag (OOB) estimates (Table S2), demonstrating robust predictive capacity.

The resulting model produced an immune-specific age for each participant. Immune age gaps (acceleration) were defined as the difference between predicted immune age and chronological age. As described previously (*23*), age gaps were further corrected by subtracting baseline gaps observed in healthy individuals and adjusting for variation across three age strata (young: ≤35 years; middle-aged: 36–60 years; older: >60 years). These corrected immune age gaps were used for all downstream analyses.

The trained models were applied to the 200HIV and 2000HIV cohorts to estimate immune-specific ages and corresponding age gaps in PWH. Immune age and immune age gap scores were then compared with systemic aging metrics derived from whole-proteome assessments—including conventional age, organ-specific ages, and organismal age—as previously described (*23*).

### DNA methylation profiling and data processing

DNA methylation profiling was performed on 1,914 samples, as previously described (*23*). Genomic DNA was extracted from EDTA-anticoagulated whole blood at the Radboudumc Genetics Department using the ChemagicStar automated system (Hamilton Robotics), which employs magnetic polyvinyl alcohol (M-PVA) bead technology. DNA concentration and purity (260/280 nm ratio) were assessed using a NanoDrop spectrophotometer. Samples were normalized to 50 ng/µL in TE buffer and randomly assigned across plates. High-quality DNA samples were processed on the Illumina Infinium MethylationEPIC BeadChip array (manifest B5).

Standard sample- and probe-level quality control (QC) procedures were applied. Raw IDAT files from the 2000HIV cohort were processed using the minfi package in R (v4.2.0)¹¹.

Samples were excluded if they exhibited sex mismatches or failed QC thresholds. Probes were removed if they had >10% missing values (detection P > 0.01), mapped to sex chromosomes, overlapped with common SNPs (minor allele frequency >5% in European populations), or aligned to multiple genomic loci. Stratified quantile normalization was applied to the remaining probes.

Methylation β-values were calculated as β = M / (M + U + 100), where M and U represent the methylated and unmethylated signal intensities, respectively.

### DNA methylation age and age gap calculation

Normalized DNA methylation β-values were used to estimate DNA methylation age (DNAm age) using four established blood-based epigenetic clocks: HorvathAge (*26*), HannumAge (*27*), PhenoAge (*28*), and GrimAge (*9*). DNAm age estimates were obtained using the online DNAm age calculator (https://dnamage.clockfoundation.org; accessed October 2024).

Preliminary age acceleration values were calculated as the difference between DNAm age and chronological age. To account for the confounding influence of chronological age, DNAm age was regressed on chronological age, and the resulting residuals—referred to as residual DNAm age gaps—were used for all downstream analyses.

### Age-related immune cell composition selection and immune cell composition age calculation

To identify immune cell subsets most strongly associated with aging in the 2000HIV cohort, we trained 500 bootstrap LASSO regression models using 356 normalized immune cell counts to predict chronological age. Thirteen immune cell populations with non-zero coefficients in all 500 models were selected. These included NK_CD56++CD16⁻, mDC, NCM_CD14⁺CD16⁺CD86⁺, CD8⁺Tc1/17, CD8⁺Tc17, mTreg_PD1⁺, CD8⁺Tcm_CD38⁺, CD8⁺Tc1_HLA-DR⁺, CD56⁺CD3⁺_CCR6⁺, CD56⁺CD3⁺_CXCR3⁺, nTreg_CXCR3⁺, Tfh_1/17_CXCR4⁺, and CD8⁺Tnaive_CXCR4⁺.

These selected immune subsets were then used to estimate immune composition–based aging in PWH within the 2000HIV cohort. Diffusion maps were generated using the destiny R package (*29*), applied to normalized frequencies of the selected immune cell populations. Diffusion pseudotime values were rescaled to a 0–1 interval and interpreted as a proxy for relative immune composition age in the PWH population.

### Linear modelling and meta-analyses

Associations between immune age, corrected immune age gap, and immune composition age were systematically evaluated in the 2000HIV cohort across a broad range of factors, including clinical characteristics, HIV disease progression markers such as latest CD4/CD8 ratio, current and cumulative antiretroviral therapy (ART) exposure, HIV reservoir size (total and intact HIV DNA in circulating CD4 cells), history of comorbidities, concomitant medication use, and immune cell composition. Linear regression models adjusted for age, sex, and ethnicity were used to assess these relationships, specified as:

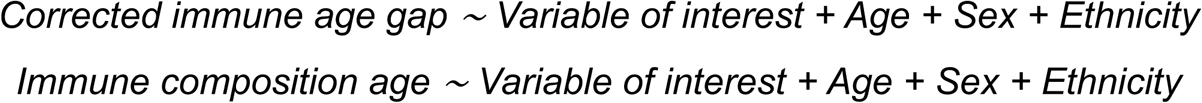

For analyses of current ART agent use, models were adjusted for age only:

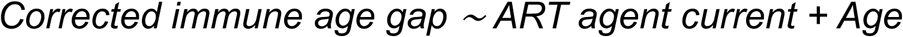

For analyses of cumulative ART agent exposure, models were adjusted for age and concomitant ART co-administration:

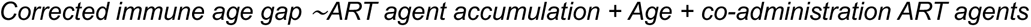

Meta-analyses based on multiple linear regression models were performed in R using the glmnet package (*30*). Where applicable, P values were adjusted for multiple testing using the Benjamini–Hochberg procedure, and results are reported as q-values or false discovery rates (FDR).

### Association analysis between immune age and heart failure markers

Associations between immune age in PWH and 26 heart failure–related protein markers, previously identified by Peterson et al., 2025 (*19*), were assessed using linear regression models of the form:

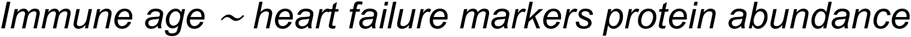

### Differential gene expression analysis and identification of age-associated genes

Bulk RNA-seq raw count data from healthy individuals (n = 423; age range: 18–92 years) were obtained from the FUSE project. Gene expression counts were normalized using the DESeq2 package (*31*). Spearman correlation coefficients were computed between normalized gene expression levels and chronological age. Genes with correlation coefficients greater than 0.3 or less than −0.3 were defined as age-associated genes.

Gene Ontology (GO) enrichment analysis was performed using the clusterProfiler package (*32*) to annotate biological pathways associated with positively and negatively age-associated genes.

Bulk RNA-seq raw count data from PWH in the 2000HIV cohort (n = 1,799; age range: 19–84 years) were processed using DESeq2. Differential gene expression analyses were conducted to compare individuals currently receiving non-nucleoside reverse transcriptase inhibitors (NNRTIs; n = 703) with those not receiving NNRTIs, and similarly for nucleoside reverse transcriptase inhibitors (NRTIs; n = 1,708 versus non-users). Chronological age was included as a covariate in all differential expression models.

## Ethics Statement

All human studies were conducted in accordance with the Declaration of Helsinki and were approved by the appropriate institutional review boards or ethics committees. Written informed consent was obtained from all participants prior to study enrollment. The clinical study was registered at ClinicalTrials.gov (NCT03994835).

## Acknowledgments

The study was supported by an unrestricted research grant from ViiV Healthcare. MGN was supported by an ERC Advanced Grant (833247) and a Spinoza Grant of the Netherlands Organization for Scientific Research.

## Author Contributions

Conceptualization: MGN, AvdV, YZ

Methodology: YZ, VM, AN, NV, MB, WV, AG, LvE, JS, ALR, IS, VK, CKB, MBe, MD

Investigation: YZ, MGN, AvdV Visualization: YZ

Funding acquisition: MGN, AvdV

Project administration: MGN, AvdV, YZ Supervision: MGN, AvdV

Writing – original draft: YZ, MGN, AvdV

Writing – review & editing: YZ, VM, AN, NV, MGN, AvdV, CR, TO, LJ, CJX, YL, LV

---

**Fig S1.**
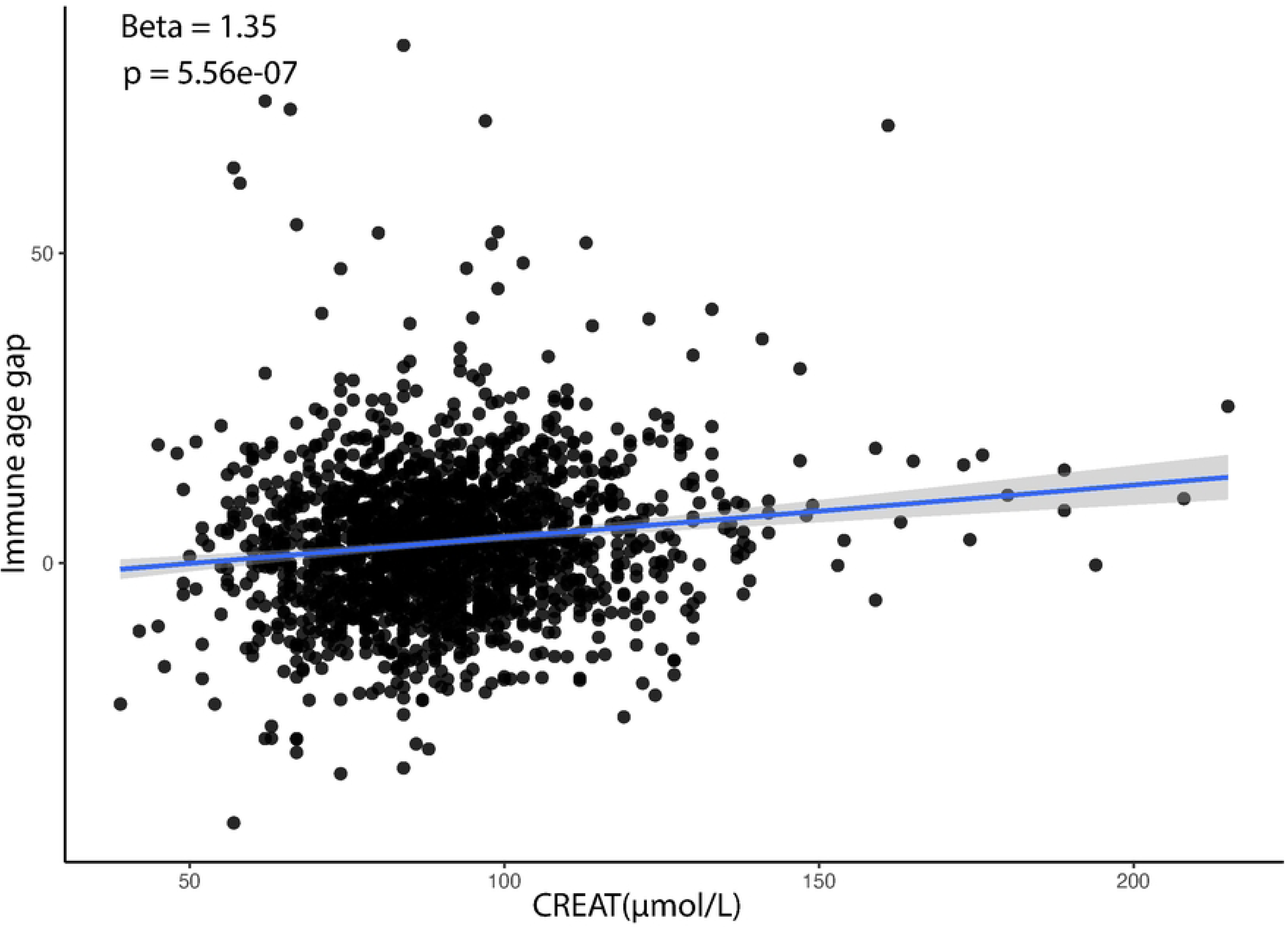
Association between immune age and serum creatinine after adjustment for dolutegravir use.

**Fig S2.**
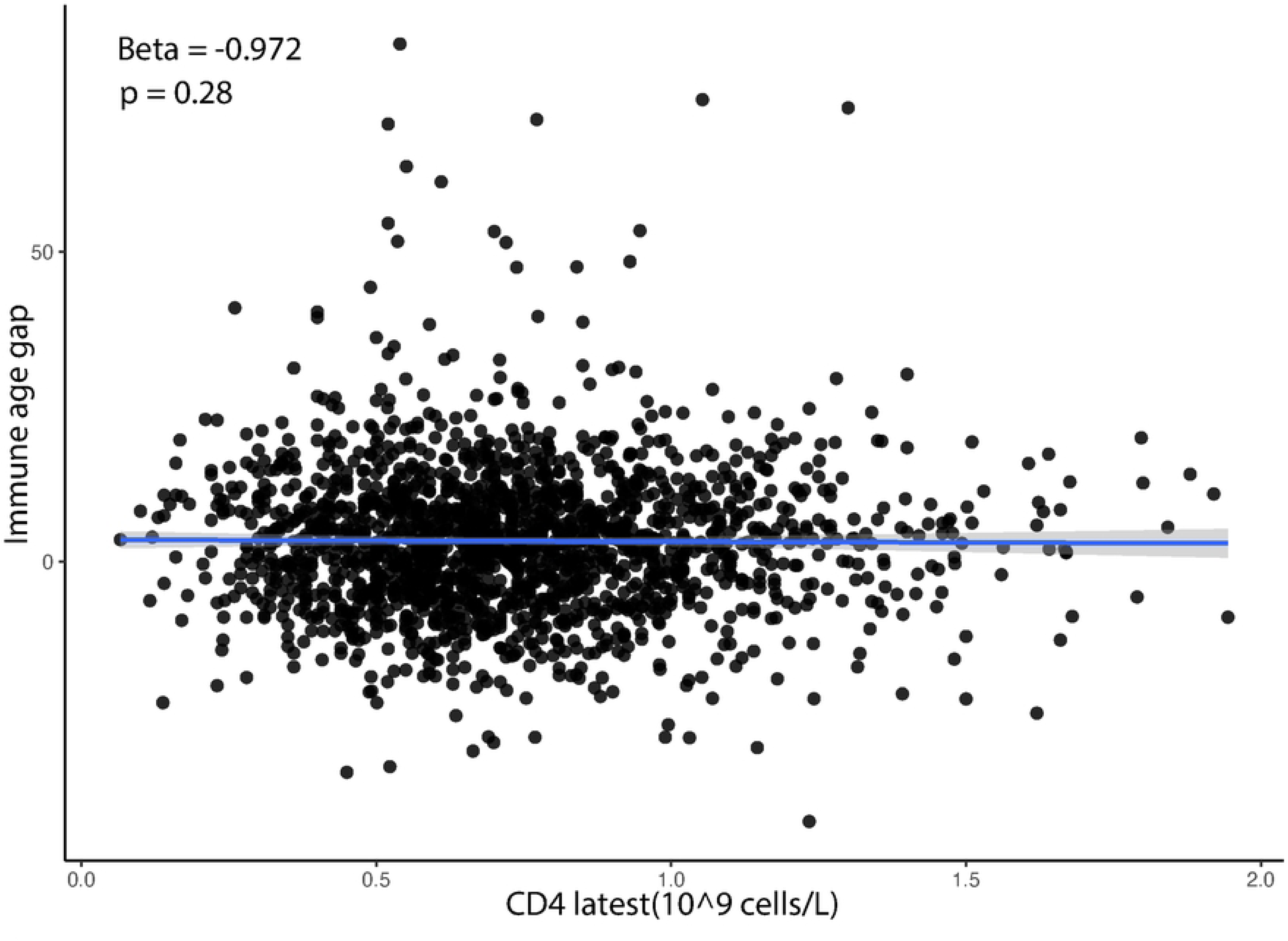
Associations between immune age and CD4⁺ T-cell count and CD4⁺/CD8⁺ T-cell ratio.

**Fig S3.**
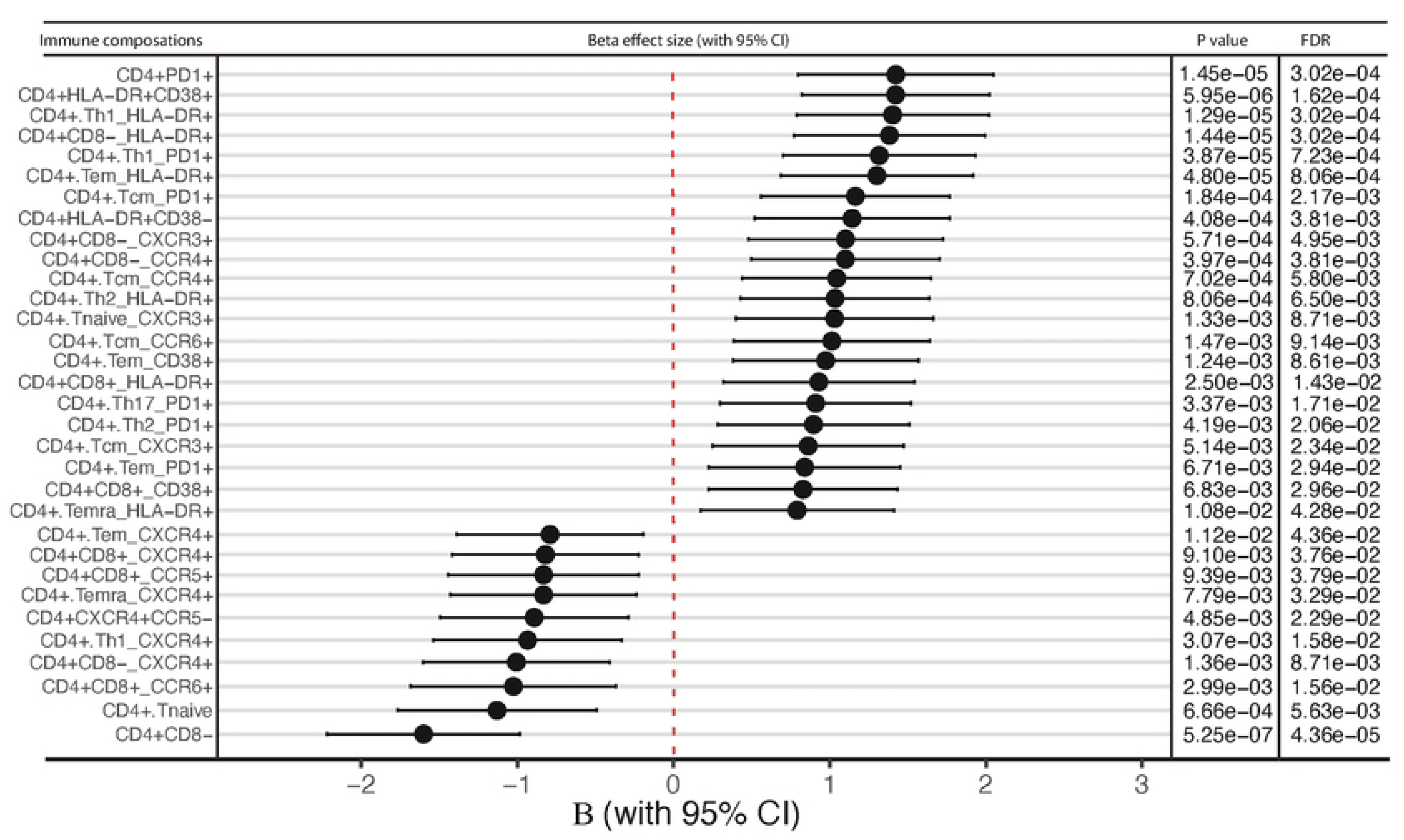
Associations between immune age and CD4⁺ T-cell subsets.

**Fig S4.**
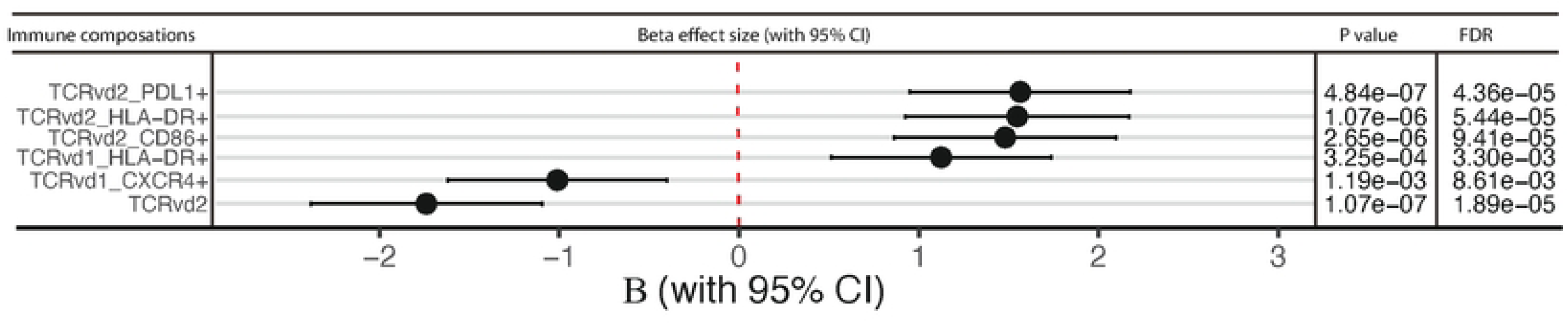
Associations between immune age and TCRγδ cell subsets.

**Fig S5.**
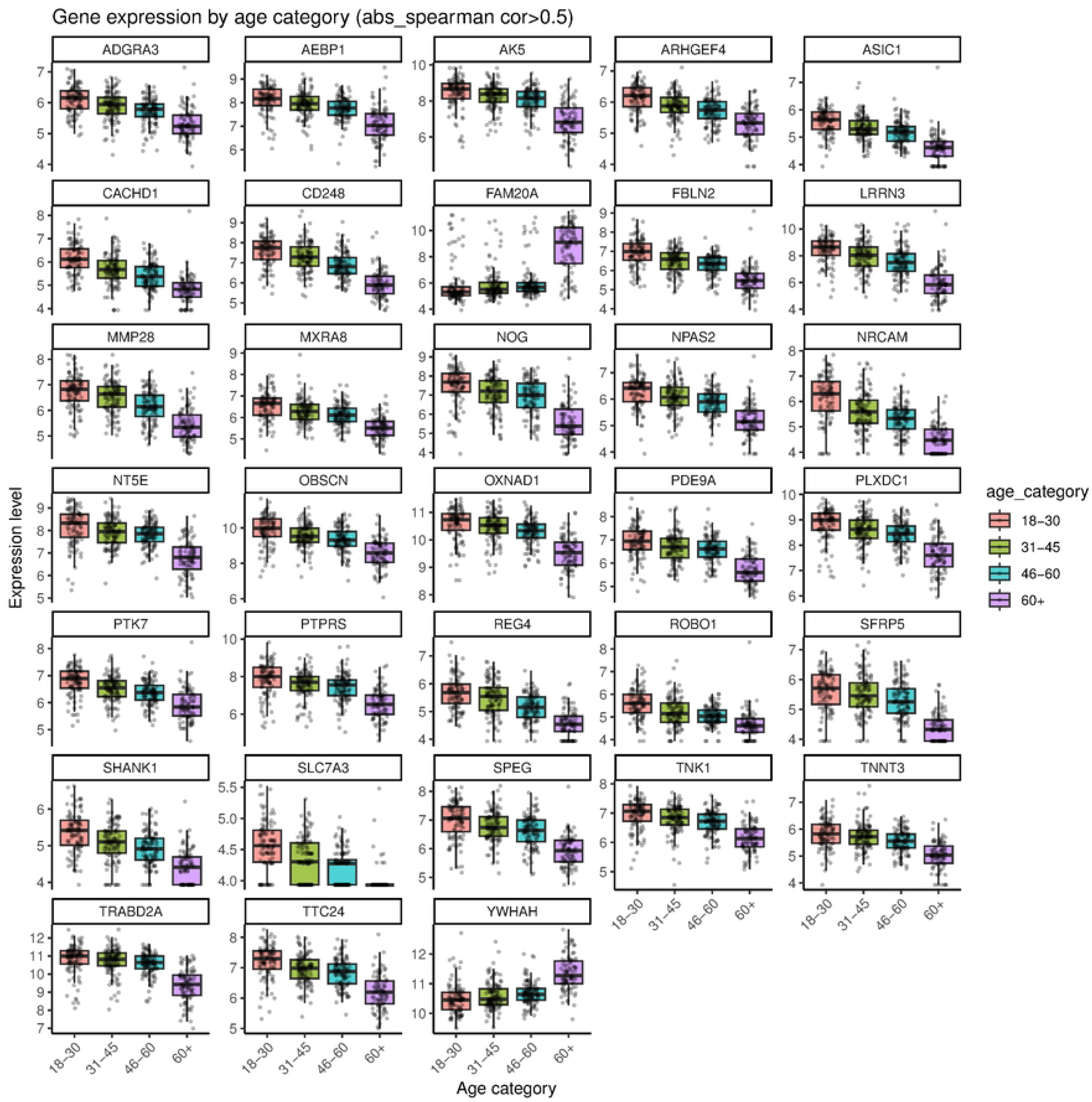
Expression patterns of strongly age-associated genes in healthy individuals.

